# Multimodal mechanisms of human centriole engagement and disengagement

**DOI:** 10.1101/2024.06.04.597290

**Authors:** Kei K Ito, Kyohei Matsuhashi, Kasuga Takumi, Kaho Nagai, Masamitsu Fukuyama, Shohei Yamamoto, Takumi Chinen, Shoji Hata, Daiju Kitagawa

## Abstract

The DNA and the centrioles are the only cellular structures that uniquely replicate to produce identical copies, which is crucial for proper chromosome segregation in mitosis. A new centriole termed “daughter” is progressively assembled adjacent to a pre-existing, “mother” centriole. Only after the daughter centriole is structurally completed as an identical copy, it disengages from its mother to become the core of a new functional centrosome. The mechanisms preventing precocious disengagement of the immature copy have been previously unknown. Here, we identify three key centriole-associated mechanisms that maintain the mother-daughter engagement: the cartwheel, the torus, and the pericentriolar material pathways. Among these, the torus is critical for establishing the characteristic orthogonal engagement between the mother and daughter centrioles. Furthermore, we show that the engagement mediated by the cartwheel and the torus pathways is released stepwise through structural changes in the daughter centriole, known as centriole blooming and centriole distancing, respectively. Disruption at any stage of these structural transitions leads to the failure of all subsequent steps, ultimately blocking centriole disengagement and centrosome conversion at the end of mitosis. Overall, this study provides a comprehensive understanding of how the maturing daughter centriole progressively disengages from its mother through multiple maturation steps, to ensure its complete structure and conversion into an independent centrosome.

## Introduction

The centrosome is a dynamic organelle that serves as the primary microtubule-organizing center (MTOC) in animal cells^1^. A cell typically contains two centrosomes throughout the cell cycle. During mitosis, the two centrosomes migrate to opposite sides of the cell to form the poles of the mitotic spindle, ensuring accurate chromosome segregation^2^. The atypical presence of three or more centrosomes results in a multipolar spindle formation and unequal chromosome segregation^3^. To maintain the correct number of centrosomes, centrosome duplication is tightly regulated and occurs only once per cell cycle^1^. Deviations from the expected proper centrosome number can lead to various diseases, including cancer and microcephaly ^4^.

The regulation of the centrosome number is primarily achieved through the control of its core component, the centriole^1^. In G1 phase, cells have two centrosomes, each composed of a matured centriole (mother centriole) surrounded by a protein matrix consisting of the torus and the pericentriolar material (PCM). The PCM is responsible for the MTOC activity of the centrosome^5^. Centrioles start to duplicate during the early S phase. The mother centriole assembles a cartwheel structure on its lateral surface, which serves as a scaffold for the formation of the daughter centriole^6^. As the cell cycle progresses, the daughter centriole undergoes an increase in width (bloom phase) followed by an increase in length (elongation phase) ^7^. During the elongation phase, the daughter centriole first acquires the A-C linker and then the inner scaffold ^7^. Further elongation occurs during mitosis, allowing the daughter centriole to reach a length similar to that of the mother^8^. After cell division, the mature daughter centriole separates from its mother (disengagement) and converts into a fully functional centrosome with its own torus and PCM (centriole-to-centrosome conversion)^9,10^. This gradual maturation process allows the daughter centriole to become an independent centrosome ^11^.

The newly-formed daughter centriole remains engaged with its mother until the end of mitosis ^1^. Precocious disengagement of the daughter centriole during mitosis results in the early centriole-to-centrosome conversion, which leads to the formation of abnormal spindles with amplified centrosomes^12,13^. When daughter centriole disengages from its mother centriole before entering mitosis, centriole reduplication occurs in addition to early conversion, which results in more severe mitotic errors^14^. Several key proteins have been identified to play crucial roles in maintaining centriole engagement. During mitosis, Cep57, a protein localizing to the mother centriole wall, is responsible for maintaining centriole engagement by stabilizing PCNT, a scaffold protein of the PCM^12,13^. Following chromosome segregation, the cleavage of PLK1-phosphorylated PCNT by the cysteine protease Separase triggers centriole disengagement^15,16,17,18^. Meanwhile, during interphase, the maintenance of centriole engagement is redundantly carried out by Cep57 and its paralog Cep57L1^14^. Although the daughter centriole undergoes a gradual maturation process involving dynamic structural alterations while attached to the mother centriole, the changes in the configuration of the mother-daughter centriole engagement from duplication to disengagement remain elusive.

In this study, we uncover the stepwise changes in the configuration and the mechanism of centriole engagement that allow the daughter centriole to disengage from its mother at the end of mitosis. Centriole engagement is initially maintained by the cartwheel structure immediately after daughter centriole formation. This engagement is then reinforced by the torus and the PCM of the mother centriole, whereby the torus plays a crucial role in establishing the characteristic orthogonal engagement between the mother and the daughter centrioles. These engagement mechanisms are sequentially released in conjunction to the structural changes of the daughter centriole during its maturation process. Interrupting the stepwise maturation of the daughter centriole at any stage halts the associated transition between engagement mechanisms, preventing the eventual disengagement and conversion of the daughter centriole into a centrosome.

## Results

### Sequential changes in the configuration of the centriole engagement during daughter centriole maturation

A daughter centriole emerges from the wall of the mother centriole in early S phase and remains engaged until the end of mitosis. While engaged with the mother centriole, the daughter gradually matures and changes its structure. To examine whether the configuration of centriole engagement also changes during this maturation process, we employed Ultrastructure Expansion Microscopy (U-ExM) to obtain high-resolution images of the centriole pairs ^19^. Most of the mother-daughter centriole pairs were engaged orthogonally in interphase (Fig. 1a, c, 80°-100°, 83.4±5.0%). However, we also observed a fraction exhibiting acute or obtuse engagement angles during this phase. As cells progressed into mitosis, the engagement angle between mother and daughter centrioles became obtuse (Fig. 1a, c, percentage of obtuse angles: 49.9±2.5%). In the subsequent G1 phase, centrioles were disengaged, which is defined as the status where distance between the CP110 signals of mother and daughter centrioles exceeds 750 nm ^14^. Additionally, the width of the daughter centriole gradually expanded concomitant with the cell cycle progression as reported recently (Fig. 1a, d, Width of daughter centrioles in late S phase: 167±26nm, in mitosis: 190±18nm, Mother centrioles in interphase: 189±19nm) ^7^. Despite the changes in engagement angle and structural width, the daughter centriole remained associated with its mother, suggesting that the configuration of centriole engagement undergoes dynamic changes throughout the process of daughter centriole maturation.

**Figure 1.**
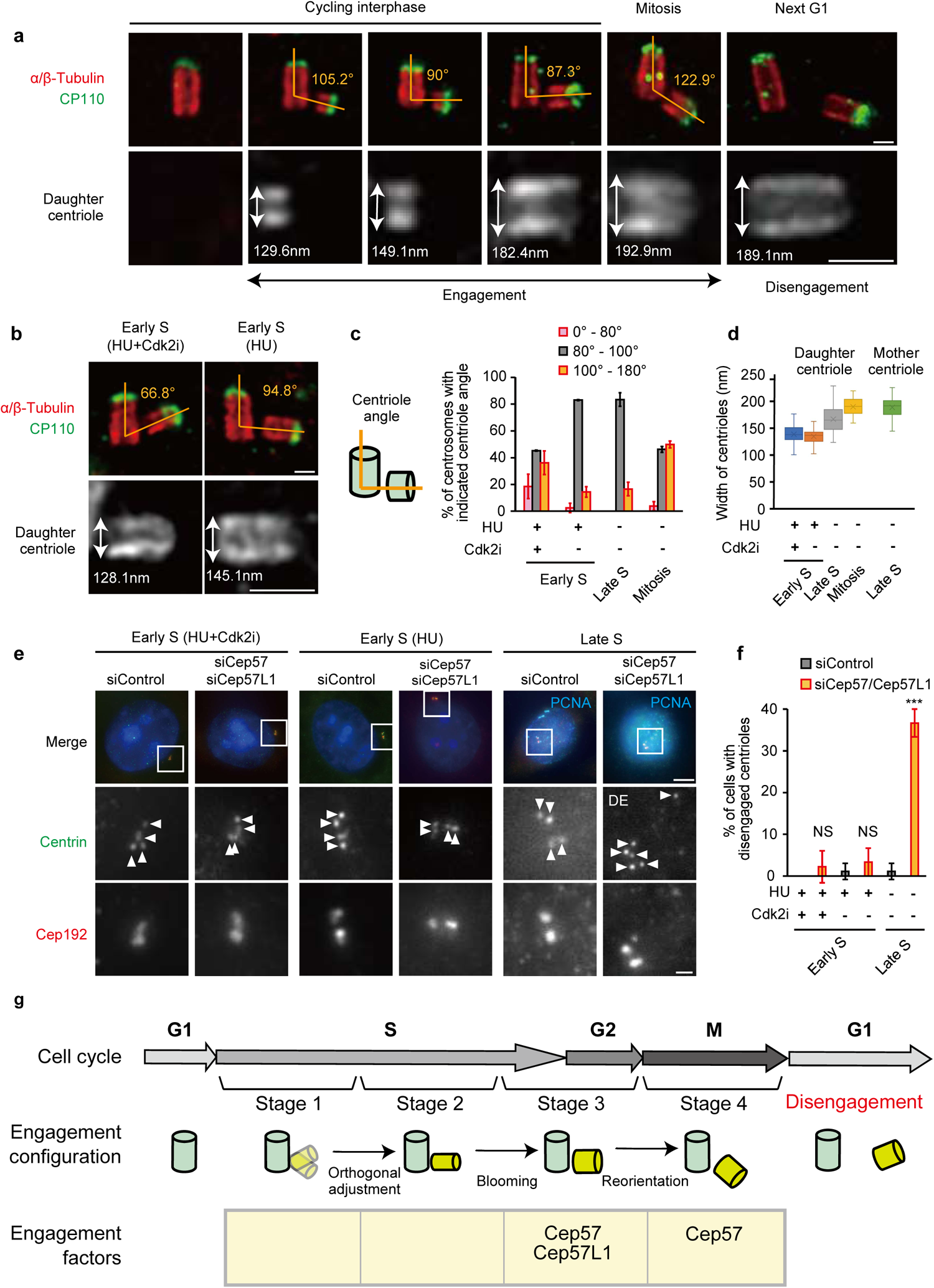
Distinct stages of centriole engagement. **(a)** Ultrastructure expansion microscopy (U-ExM) images of centrioles through the cell cycle in HeLa cells. Scale bars: 200 nm. **(b)** U-ExM images of centrioles in HeLa cells treated with hydroxyurea (HU) or HU + Cdk2 inhibitor III (Cdk2i). Scale bars: 200 nm. **(c)** Quantification of the frequency of centrosomes with indicated mother-daughter centriole angles (0-80°, 80-100°, and 100-180°) observed by STED microscopy. n = 3 independent experiments, 20 centrosomes each. **(d)** Quantification of diameters of daughter centrioles observed by STED microscopy. n = 40 centrioles pooled from 3 independent experiments. **(e)** Representative immunofluorescence images of HeLa cells transfected with siControl or siCep57/Cep57L1 under DDW, HU or HU + Cdk2i treatment. Scale bars: 5 µm, 1 µm. **(f)** Quantification of the frequency of cells with disengaged centrioles in (e). n = 3 independent experiments, 50 cells each. P values were calculated by two-tailed unpaired Welch’s t-test with Bonferroni correction. **(g)** Schematic model of the four stages of centriole engagement throughout the cell cycle. Stage 1: flexible intercentriolar link and narrow daughter centrioles. Stage 2: orthogonal intercentriolar link and narrow daughter centrioles. Stage 3: expanded daughter centrioles, engaged to the mother by Cep57-Cep57L1. Stage 4: daughter centrioles at obtuse angle, engaged to the mother solely by Cep57. Centrioles disengage in the following G1 phase. Data are represented as mean ± s.d.

Daughter centrioles that engaged at non-orthogonal angles (<80° or >100°) appear thinner in width (Fig. 1a), suggesting an incipient configuration of centriole engagement immediately after duplication. To confirm this, we arrested the cell cycle at early S phase using Hydroxyurea (HU, a DNA replication inhibitor) and analyzed the centriole pairs using U-ExM (Fig. 1b). Under HU treatment, daughter centrioles appeared thinner (Fig. 1b, d, 135±13nm, mean ± s.d.), but their engagement angle with the mother centriole was orthogonal (Fig. 1b, c). This suggests that the non-orthogonal configuration may occur at an earlier timepoint, before the HU treatment conditions. To test this hypothesis, we treated cells with a Cdk2 inhibitor simultaneously with HU, as Cdk2 is thought to regulate the cell cycle progression from late G1 to S phase and was shown to be involved in centriole/centrosome formation^20–23^. After Cdk2 inhibition, we observed a decrease in the width of the daughter centrioles (Fig. 1b, d: 139±18nm), together with a larger fraction of centrioles that engaged with their mother at acute or obtuse angles (Fig. 1b, c). These results suggest that the configuration of the mother-daughter centriole engagement is flexible immediately after duplication, transitioning to a stable, orthogonal orientation as the cell cycle progresses.

These results further reveal that there are at least four distinct configurations of mother-daughter centriole engagement (Fig. 1g). Stage 1 is characterized by the thin structure of the daughter centriole and the flexible association to the mother that is observed immediately after centriole duplication. Stage 2 occurs in early S phase, in which the width of the daughter centriole remains thin but the link to the mother adopts a more stable orthogonal configuration. During late S and G2 phases the centriolar configuration enters Stage 3, which is marked by the increase in thickness of the daughter centriole and the orthogonal mother-daughter association. Lastly, in Stage 4 that takes place during mitosis, the majority of the centriole pairs are engaged under an obtuse angle. The transient configurations of the first two stages can be specifically enriched by treating cells with HU+Cdk2i (Stage 1) or HU alone (Stage 2), while untreated cells also display late S/G2 (Stage 3) or mitotic (Stage 4) engagement profiles.

### The mechanism of the centriole engagement is diverse in interphase

We previously reported that suppressing the expression of both Cep57 and Cep57L1 causes precocious centriole disengagement in interphase frequently followed by centriole re-duplication (Fig. s1a, s1b)^14^, which was confirmed using knockout cells (Fig. s2a-e). To investigate whether all three configurations of centriole engagement found in interphase (Stages 1-3) are maintained by Cep57 and Cep57L1, we knocked-down their expression and assessed the centrioles under each condition. Suppressing Cep57 and Cep57L1 caused centriole disengagement and reduplication in cycling late S phase (Stage 3) cells, but surprisingly, not in early S cells treated with HU + Cdk2 inhibitor (Stage 1) or HU alone (Stage 2) (Fig. 1e, f). Consistent with this result, Cep57 and Cep57L1 co-depletion did not cause centriole disengagement in cycling early S phase (Stage 1 or 2) (Fig. s2f, g). Therefore, early S phase centriole engagement (Stage 1, 2) is maintained through unknown mechanisms, distinct from the Cep57- and Cep57L1-dependent pathway in late S and G2 phase (Stage 3). Furthermore, previous reports demonstrated that suppressing Cep57 alone leads to centriole disengagement in mitosis (Stage 4, Fig. s1a, c) ^12,24^. Taken together, these results suggest that there are multiple mechanisms in place to maintain centriole engagement in the different phases of the cell cycle (Fig. 1g).

### The cartwheel structure contributes to centriole engagement in early S phase: Stage 1 and Stage 2

Since Stage 1 in early S phase occurs immediately after the onset of centriole duplication, we hypothesized that the cartwheel, a key structure involved in this process, might play a role in centriole engagement in this stage. This hypothesis is supported by a previous report that demonstrated the involvement of the cartwheel in maintaining centriole engagement during interphase ^25^. To test the hypothesis, we removed the cartwheel after daughter centriole formation using the PLK4 inhibitor centrinone ^26^. Cells were first arrested in early S phase by treatment with HU+Cdk2 inhibitor for 24 hours, inducing Stage 1 state. At this point, all cells displayed newly-formed daughter centrioles with cartwheels, associated to their respective mothers within centrosomes (Fig. 2a, b, HU+Cdk2i: 100.0%). Upon further addition of PLK4 inhibitor for 24 hours, the cartwheel was lost in approximately half of the cells (44.4 ±15.0%). Unexpectedly, in Stage 1 cells that had lost the cartwheel, the number of centrioles decreased from four to two in about 90% of cells (Fig. 2a, b, 4 centrioles: 11.1 ± 11.1%), suggesting a defect in the integrity of the daughter centriole. Interestingly, among the cells with four remaining centrioles after cartwheel loss, approximately 10% exhibited centriole disengagement (Fig. s2h). We therefore assumed that daughter centrioles that disengage from the mother centriole at this timepoint, are unstable and undergo rapid disintegration.

**Figure 2.**
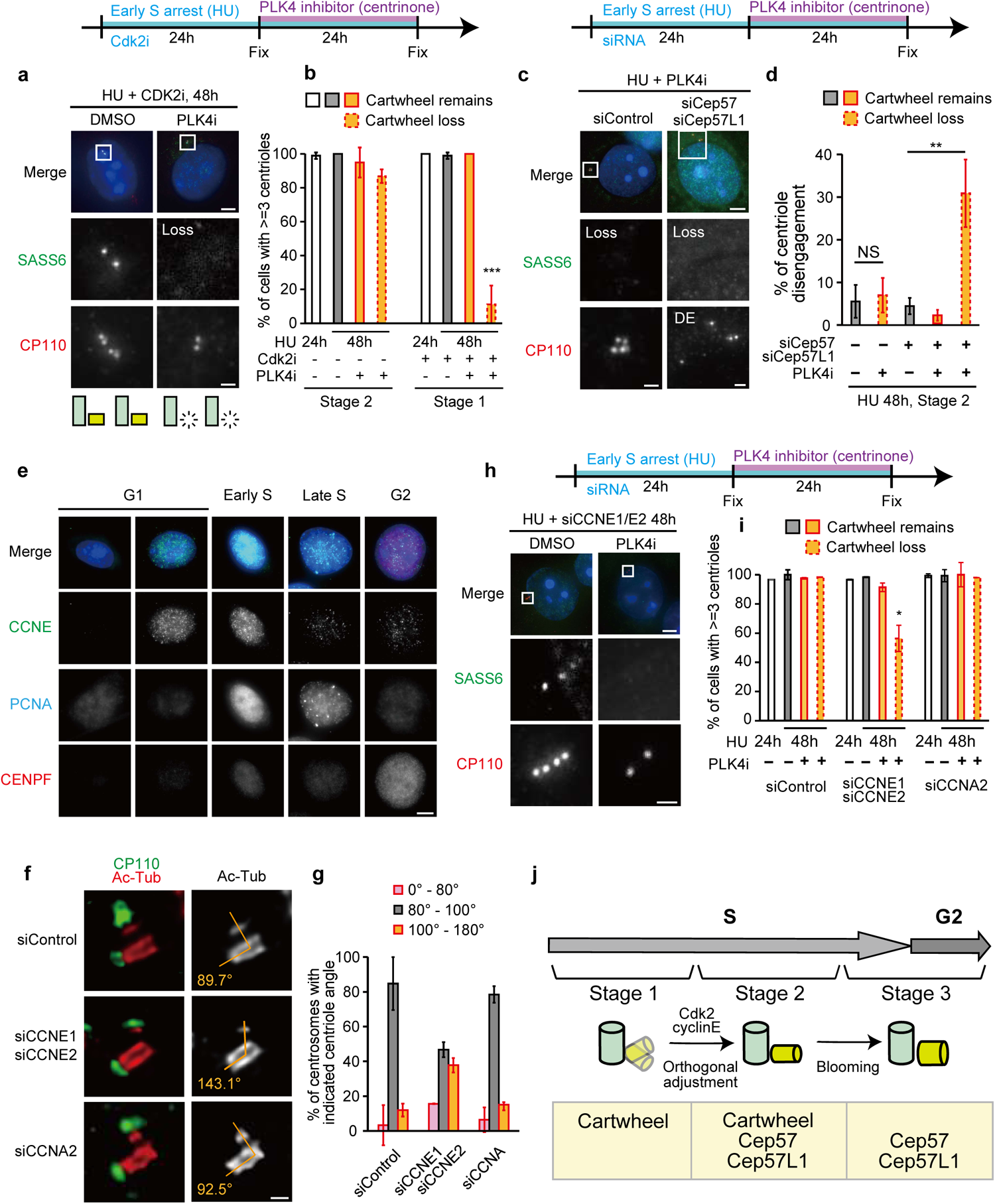
The cartwheel contributes to centriole engagement in early S phase. **(a)** Representative immunofluorescence images of HeLa cells subjected to a cartwheel removal assay. HeLa cells were treated with HU and Cdk2 inhibitor III (Cdk2i) for 24 hours, followed by treatment with a PLK4 inhibitor for another 24 hours. Scale bars: 5 µm, 1 µm. **(b)** Quantification of the frequency of cells containing three or more centrioles in (a). Solid bars indicate the centrosomes that retain the cartwheel structure, while dashed bars represent the centrosomes that have lost the cartwheel structure. n = 3 independent experiments, 30 cells each. P values were calculated by Dunnett’s multiple comparisons test. **(c)** Representative immunofluorescence images of HeLa cells subjected to a cartwheel removal assay. HeLa cells were treated with HU and transfected with indicated siRNAs for 24 hours, followed by treatment with a PLK4 inhibitor for another 24 hours. Scale bars: 5 µm, 1 µm. **(d)** Quantification of the frequency of cells with disengaged centrioles in (c). n = 3 independent experiments, 30 cells each. P values were calculated by two-tailed unpaired Welch’s t-test (siControl) and Dunnett’s multiple comparisons test (siCep57 + siCep57L1). **(e)** Immunofluorescence images of HeLa cells throughout the cell cycle. The cell cycle phase was determined by the signal pattern of PCNA and CENP-F. Scale bar: 5 µm. **(f)** Representative STED microscopy images of centrosomes in HeLa cells in late S phase transfected with indicated siRNAs. Scale bar: 200 nm. **(g)** Quantification of the frequency of centrosomes with the indicated mother-daughter centriole angles in (f). n = 3 independent experiments, 20 centrosomes each. **(h)** Representative immunofluorescence images of HeLa cells subjected to a cartwheel removal assay. HeLa cells were treated with HU and siRNA for 24 hours, followed by treatment with a PLK4 inhibitor for another 24 hours. Scale bars: 5 µm, 1 µm. **(i)** Quantification of the frequency of cells containing three or more centrioles in (h). n = 3 independent experiments, 30 cells each. P values was calculated by Dunnett’s multiple comparisons test. **(j)** Schematic model of the mechanisms and configurations of centriole engagement during interphase. Data are represented as mean ± s.d.

In contrast, Stage 2 cells retained four centrioles even when the cartwheel was lost, and the engagement between mother and daughter centrioles was still maintained, indicating that cartwheel loss does not affect engagement stability (Fig. 1b, 2c, d, Disengagement: 4.4 ± 1.9%). We hypothesized that Cep57 and Cep57L1, which are involved in maintaining centriole engagement in Stage 3, might also work redundantly with the cartwheel in maintaining centriole engagement in Stage 2. To test this hypothesis, we removed the cartwheel and simultaneously suppressed Cep57 and Cep57L1 expression in Stage 2 cells (Fig. 2c, d). Strikingly, precocious centriole disengagement was observed in this condition (Fig. 2c, d, 30.9 ± 7.9%). In the cells retaining the cartwheel after PLK4 inhibitor treatment, we did not observe precocious centriole disengagement, indicating that it is not PLK4 but the cartwheel that maintains engagement (Fig. 2d, 3.3 ± 3.3%). These findings indicate that centriole engagement in Stage 2 is redundantly maintained by Cep57, Cep57L1, and the cartwheel.

### Orthogonal centriole engagement is established by CDK2-cyclin E: from Stage 1 to Stage 2

Inhibiting Cdk2 results in an accumulation of centrioles at Stage 1 of centriole engagement, implying that its activity is required for the transition from Stage 1 to Stage 2. CDK2 is activated by forming complexes with either cyclin E or cyclin A (Fig. 2e, 3a)^27^. To elucidate the specific cyclin contributing to the configuration and mechanism changes behind the transition from Stage 1 to Stage 2, we knocked down each cyclin and observed centrioles during S phase. Control cells showed orthogonal centrioles, while depletion of cyclin E but not cyclin A displayed unstable centriole angles (Fig. 2f, g, 80°-100°, siControl: 84.7 ± 4.7%, siCCNE1/E2: 46.7 ± 4.7%, siCCNA2: 78.4 ± 8.0%), suggesting that CDK2-cyclin E alters the centriole configuration from Stage 1 to Stage 2. The mechanism of centriole engagement also differs between Stage 1 and Stage 2, particularly in the dependency on the cartwheel structure. Similar to the CDK2 inhibitor treatment, cartwheel removal by PLK4 inhibitor caused centriole disengagement and loss of daughter centrioles in cells with suppressed cyclin E expression (Fig. 2h, i, 4 centrioles: 56.4 ± 8.9%). These findings suggest that CDK2-cyclin E mediates the transition of both the configuration and the mechanism of centriole engagement from Stage 1 to Stage 2 (Fig. 2j).

**Figure 3.**
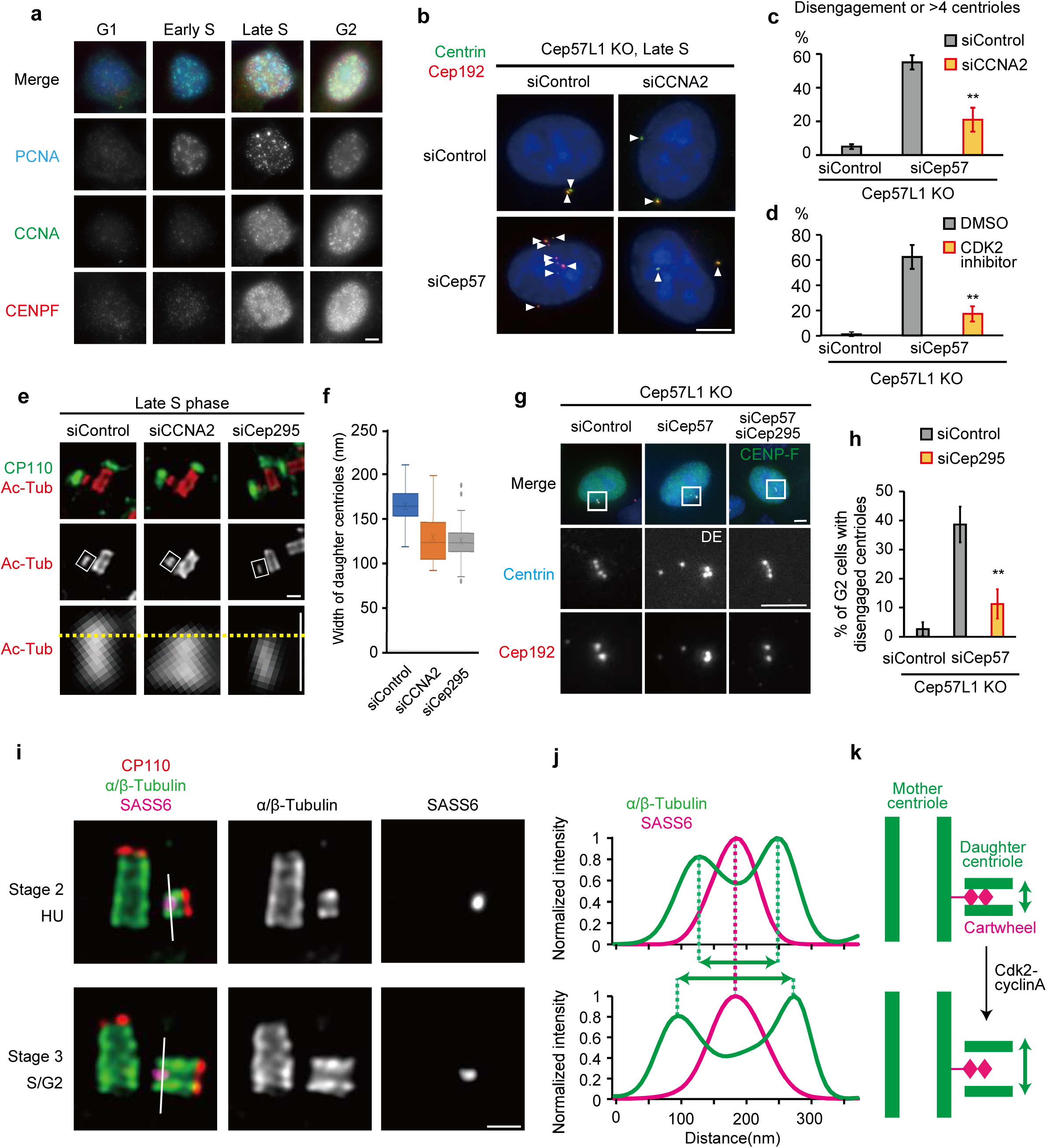
Daughter centriole blooming releases the cartwheel centriole engagement pathway. **(a)** Immunofluorescence images of HeLa cells throughout the cell cycle. The cell cycle phase was determined by the signal pattern of PCNA and CENP-F. Scale bar: 5 µm. **(b)** Representative immunofluorescence images of HeLa Cep57L1KO cells transfected with siControl and siCep57, in the presence or absence of siCCNA2, Scale bar: 5 µm. **(c)** Quantification of the frequency of cells with disengaged centrioles in (b). The cell cycle phase was determined by the signal pattern of PCNA. n = 3 independent experiments, 30 cells each. P value was calculated by two-tailed unpaired Welch’s t-test with Bonferroni correction. **(d)** Quantification of the frequency of cells with disengaged centrioles in late S phase of Cep57L1KO cells transfected with siControl and siCep57, in the presence or absence of Cdk2 inhibitor. The cell cycle phase was determined by the signal pattern of PCNA. n = 3 independent experiments, 30 cells each. P value was calculated by two-tailed unpaired Welch’s t-test with Bonferroni correction. **(e)** Representative STED microscopy images of HeLa cells transfected with siControl, siCCNA2, or siCep295. Scale bar: 200 nm. **(f)** Quantification of the width of the daughter centrioles. n = 40 centrioles pooled from 3 independent experiments. **(g)** Representative immunofluorescence images of HeLa Cep57L1 KO cells transfected with siControl, siCep57, or siCep57 + siCep295. Scale bars: 5 µm, 1 µm. **(h)** Quantification of the frequency of cells with disengaged centrioles in (g). n = 3 independent experiments, 30 cells each. P value was calculated by two-tailed unpaired Welch’s t-test with Bonferroni correction. **(i)** U-ExM images of centrioles in HeLa cells treated with HU (early S) or DDW (late S or G2). Scale bar: 200 nm. **(j)** Quantification of the signal intensity of α/β-tubulin and SASS6 in (i). **(k)** Schematic model for the release of the cartwheel engagement by daughter centriole blooming. Data are represented as mean ± s.d.

### Centriole blooming leads to a decreased reliance on the cartwheel for maintaining centriole engagement: from Stage 2 to Stage 3

In late S phase, the configuration and the mechanism of centriole engagement changes: daughter centrioles bloom and the cartwheel becomes dispensable for maintaining the engagement(Fig. 2j, Stage 2→3). In this phase, cyclin A, which forms a complex with CDK2 ^27^, begins to accumulate (Fig. 3a). We therefore examined whether the CDK2-cyclin A complex causes the transition from Stage 2 to Stage 3. In Cep57L1 knockout cells, Cep57 depletion led to precocious centriole disengagement in late S phase, whereas simultaneous depletion of cyclin A reduced this phenotype (Fig. 3b, c, siCep57: 55.0 ± 4.2%, siCep57+siCCNA2: 21.1% ± 7.1%). Furthermore, CDK2 inhibitors also suppressed the precocious centriole disengagement in the absence of Cep57 and Cep57L1 (Fig. 3d, DMSO: 62.7±9.5%, CDK2i: 17.3±5.7%). STED microscopy analysis revealed that depleting cyclin A resulted in daughter centrioles that remained as thin as those observed in early S phase (Fig. 3e, f, siControl: 165 ± 20 nm, siCCNA2: 127 ± 22 nm), indicating lack of daughter centriole blooming. These findings suggest that CDK2-cyclin A mediates the transition of both the configuration and the mechanism of centriole engagement from Stage 2 to Stage 3.

To investigate the relationship between daughter centriole blooming and the changes in the engagement mechanism, we sought to suppress the former by depletion of Cep295, which plays a role in the maturation process of the daughter centriole ^28,29^. Indeed, employing this approach resulted in thin daughter centrioles even in late S phase (Fig. 3e, f, siCep295: 126±24nm). When their blooming was suppressed by Cep295 depletion, the daughter centrioles were significantly less likely to disengage in Cep57-depleted Cep57L1 knockout cells (Fig. 3g, h, siCep57: 38.7±11.3%, siCep57+siCep295: 11.3±5.0%). These results suggest that the daughter centriole blooming causes the transition of the centriole engagement mechanism from Stage 2 to Stage 3. After this transition, the cartwheel is no longer required for centriole engagement. U-ExM analysis revealed that the distance between the cartwheel and the microtubule wall expanded during daughter centriole blooming (Fig. 3i, j). Thus, we speculate that the daughter centriole blooming weakens the connection between the daughter centriole wall and its internal cartwheel (Fig. 3k).

### Centriole engagement from late S phase to G2 phase is maintained by the torus and the PCM around mother centrioles: Stage 3

In Stage 3 spanning from late S phase to G2 phase, the localization of Cep57 and Cep57L1 around the wall of the mother centriole is necessary for centriole engagement (Fig. s1) ^14^. However, STED microscopy revealed that there is a distance between the outer edge of the Cep57/Cep57L1 ring (radius: 135 nm) and the proximal end of the daughter centriole (approximately 160 nm away from the center of the mother centriole), suggesting the presence of other factors that link them (Fig. 4a, b). To identify these factors, we examined the functions of Cep63, Cep152, and PCNT, which are known to interact with Cep57/Cep57L1 and are localized further away from the center of the mother centriole (Fig. 4a, b) ^12,30–32^,^33^. However, single siRNA treatment against these proteins did not result in precocious centriole disengagement in G2 phase (Fig. 4d). Considering the possibility that centriole engagement is maintained in a complementary manner, we then performed a co-depletion screening of Cep57, Cep57L1, Cep63, Cep152, and PCNT. Among all the combinations, precocious centriole disengagement was observed in only three combinations: siCep57 + siCep57L1, siCep57 + siCep63, and siCep63 + siPCNT (Fig. 4c, d, siControl: 2.0±2.7%, siCep57+siCep57L1: 64.3±0.5%, siCep57+siCep63: 47.2±2.6%, siCep63+siPCNT: 33.7±3.3%). PCNT is known to maintain centriole engagement during mitosis ^17,34,16^, while the role of Cep63 has not been reported. To validate the involvement of Cep63 in centriole engagement, we generated a Cep63 knockout cell line using CRISPR-Cas9 (Fig. s3a-c). Although previous studies have reported multipolar spindle formation and a reduction in centriole number in Cep63 knockout cells ^35,36^, these phenotypes were not observed in the cell line generated in this study (Fig. s3b, d, e). When Cep63 knockout cells were treated with siCep57 or siPCNT, precocious centriole disengagement was observed in interphase, consistent with the results of the co-depletion (Fig. 4c, d, Fig. s3f, g, siControl: 2.2±1.9%, siCep57: 41.2±8.4%, siPCNT: 35.9±3.0%). These results indicate that Cep63 and PCNT are involved in centriole engagement from late S to G2 phase (Stage 3), in addition to Cep57 and Cep57L1.

**Figure 4.**
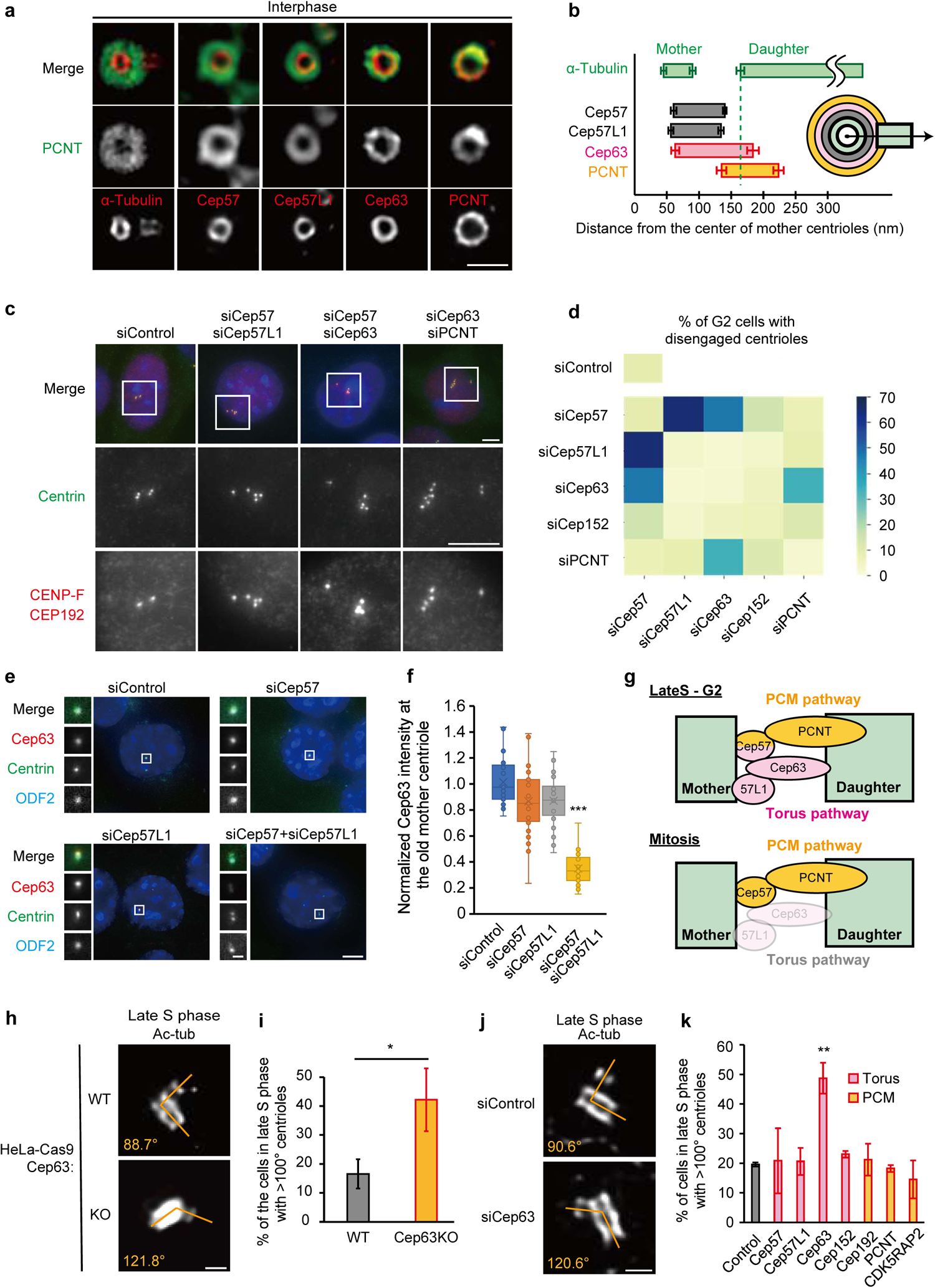
Centriole engagement is maintained redundantly by the torus and the PCM pathways from late S to G2 phase. **(a)** Representative STED microscopy images of centrioles and surrounding centrosomal proteins in HeLa cells. Scale bar: 200 nm. **(b)** Quantification of the locations of the indicated centrosomal proteins relative to the center of the mother centriole. The end of the signal was defined as the location where the signal intensity dropped to 20% of its peak value. n = 30 centrioles pooled from 3 independent experiments. **(c)** Representative immunofluorescence images of HeLa cells transfected with the indicated siRNAs. Scale bars: 5 μm. **(d)** Quantification of the frequency of cells with disengaged centrioles in (c). n = 3 independent experiments, 50 cells each. **(e)** Representative immunofluorescence images of HeLa cells transfected with the indicated siRNAs. Scale bars: 5 μm, 1 µm. **(f)** Quantification of signal intensity of Cep63 at the old mother centriole in (e). n = 50 centrioles. P value was calculated by Dunnett’s multiple comparisons test. **(g)** Schematic model for the molecular mechanisms of centriole engagement. **(h)** Representative STED microscopy images of centrioles in late S phase of HeLa-Cas9 and Cep63KO HeLa-Cas9 cells. Scale bar: 200 nm. **(i)** Quantification of the frequency of the centrosomes, in which the centrioles are connected at obtuse angle in (h). n = 3 independent experiments, 20 cells each. P value was calculated by two-tailed unpaired Welch’s t-test. **(j)** Representative STED microscopy images of centrioles in late S phase upon indicated siRNA treatments. Scale bar: 200 nm. **(k)** Quantification of the frequency of the centrosomes, in which the centrioles are connected at obtuse angle in (j). n = 3 independent experiments, 20 cells each. P values was calculated by Dunnett’s multiple comparisons test. Data are represented as mean ± s.d.

While PCNT is also known to maintain centriole engagement during mitosis with Cep57 ^12^, Cep63 appears to be an interphase-specific factor along with Cep57L1, as knocking out Cep63 does not cause centriole disengagement in mitosis (Fig. s3h). Indeed, simultaneous depletion of Cep63 and PCNT, or Cep63 and Cep57 caused centriole disengagement, suggesting that interphase engagement is supported by two pathways: the interphase-specific Cep63/Cep57L1 pathway and the Cep57/PCNT pathway, which also functions in mitosis. In interphase, Cep63 and Cep57L1 interact with each other and are localized in close proximity, whereby the Cep63 ring surrounds Cep57L1 from the outside on the mother centriole (Fig. 4a, s3i) ^31^. Therefore, we thought that the role of Cep57L1 in the engagement may be to recruit Cep63. However, Cep57L1 depletion did not affect the amount of Cep63 at the centrosome (Fig. 4e, f). Interestingly, when Cep57 and Cep57L1 were simultaneously depleted, Cep63 levels were significantly reduced (Fig. 4e, f)^14,31^. Thus, in the interphase-specific pathway, Cep57 and Cep57L1 redundantly control the engagement through Cep63. This finding is consistent with the observation that simultaneous depletion of either Cep57 + PCNT or Cep57L1 + PCNT did not lead to premature centriole disengagement, as the remaining protein (Cep57L1 or Cep57, respectively) could still maintain engagement through Cep63. Therefore, from late S to G2 phase (Stage 3), centriole engagement is maintained by two pathways: the Cep57-PCNT pathway and the Cep57-Cep57L1-Cep63 pathway. Since the outer proteins in each pathway, PCNT and Cep63, are components of the PCM^5^ and the torus^32^, respectively, we define the Cep57-PCNT as the PCM pathway and the Cep57-Cep57L1-Cep63 as the torus pathway (Fig. 4g).

### The torus protein Cep63 ensures the orthogonal engagement of mother-daughter centrioles

Remarkably, the orthogonal engagement of mother-daughter centrioles during interphase was compromised in Cep63 knockout cells (Fig. 4h, i, obtuse centriole angle, HeLa-Cas9: 16.6±5.0%, Cep63KO: 42.2±10.8%). The obtuse centriole angle is normally observed in mitosis (Stage 4), suggesting that the torus pathway loses its function in centriole engagement during the transition from Stage 3 to Stage 4. Depletion of other torus or PCM proteins, which does not disrupt the torus engagement pathway, did not result in this obtuse centriole angle (Fig. 4j, k). Therefore, the torus engagement pathway appears to be essential for maintaining the orthogonal engagement of mother-daughter centriole pairs.

### Centriole distancing mediated by the loss of the daughter’s proximal end drives the transition of the centriole engagement mechanism: from Stage 3 to Stage 4

The obtuse configuration observed in early mitosis was reported to be caused by centriole distancing, whereby the distance between the wall of the mother centriole and the proximal end of the daughter centriole slightly increases ^37^. This phenomenon is mediated by the activity of the kinase PLK1 (Fig. 5a, b) ^37^. We hypothesized that the centriole distancing also altered the mechanism of centriole engagement from Stage 3 to 4. In mitosis, when the PCM pathway was inhibited by suppressing Cep57 or PCNT expression, centrioles disengaged precociously (Fig 5c, d). However, PLK1 inhibition prevented both centriole distancing and precocious disengagement (Fig. 5a, b, c, d). Furthermore, overexpression of constitutive active PLK1, which induces precocious centriole distancing (Fig. s4a, b)^37^, caused precocious centriole disengagement upon either Cep57 or PCNT depletion in interphase (Fig. s4c, d). These results imply that centriole distancing is necessary and sufficient for the change in engagement mechanism.

**Figure 5.**
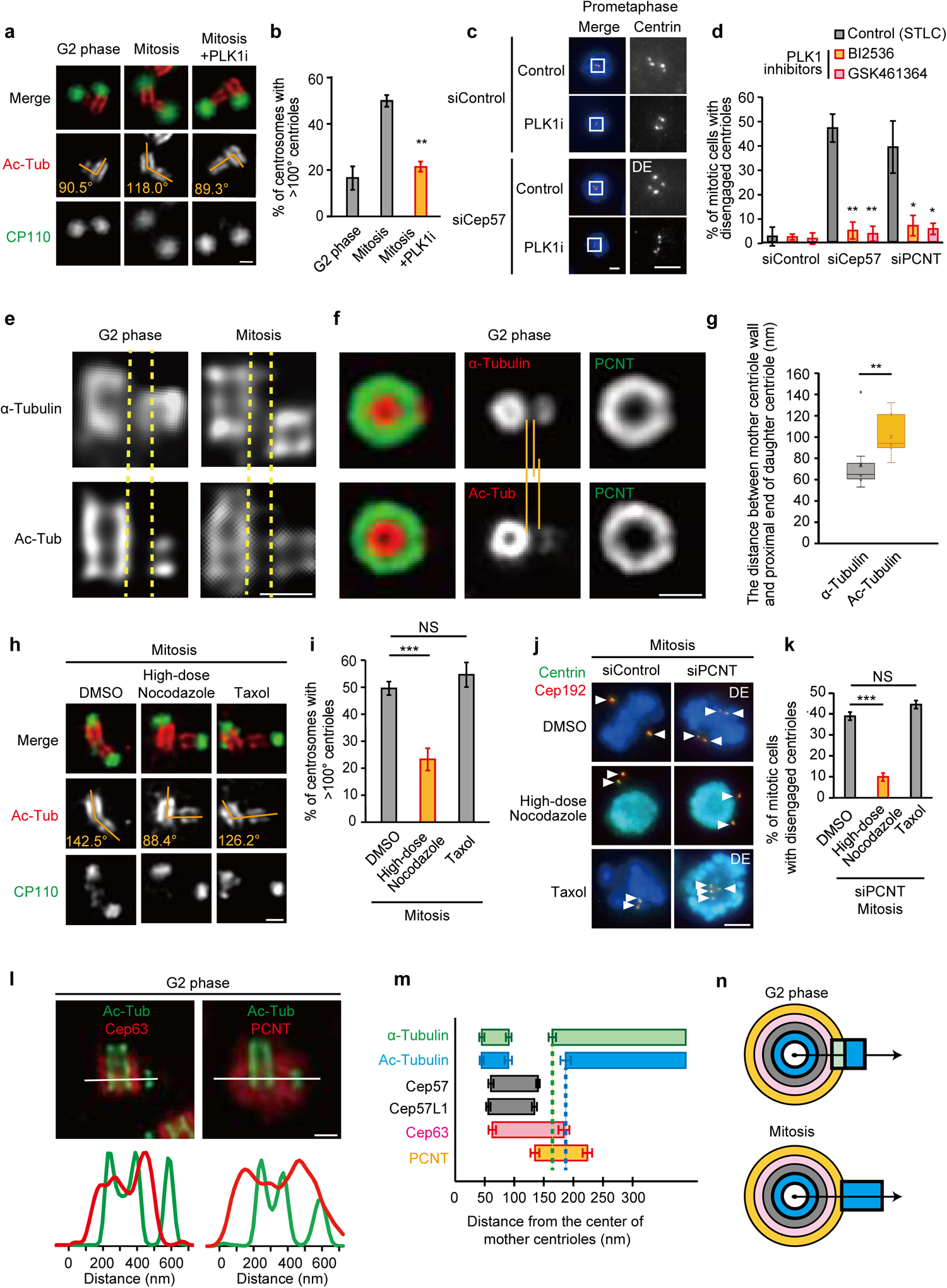
Centriole distancing releases centriole engagement by the torus pathway. **(a)** Representative STED microscopy images of centrioles in G2 phase and mitosis treated with PLK1 inhibitor. Scale bar: 200 nm. **(b)** Quantification of the frequency of the centrosomes, in which the centrioles are connected at obtuse angle in (h). n = 40 centrioles pooled from 3 independent experiments. P value was calculated by two-tailed unpaired Welch’s t-test with Bonferroni correction. **(c)** Representative immunofluorescence images of HeLa cells in mitosis transfected with siControl and siCep57, in the presence or absence of PLK1 inhibitor (BI2536). Scale bar: 5 µm. **(d)** Quantification of the frequency of mitotic cells with disengaged centrioles in (c). n = 3 independent experiments, 30 cells each. P values were calculated by Dunnett’s multiple comparisons test. **(e)** STED microscopy images of centrioles in G2 phase and mitosis immunostained with anti-α-tubulin or anti-acetylated tubulin antibodies. **(f)** Representative STED microscopy images of centrioles in G2 phase immunostained with anti-PCNT and anti-α-tubulin or anti-acetylated tubulin antibodies. Scale bar: 200 nm. **(g)** Quantification of the distance between the mother centriole wall and the proximal end of the daughter centriole in (f). n = 40 centrioles pooled from 3 independent experiments. P value was calculated by two-tailed unpaired Welch’s t-test. **(h)** Representative STED microscopy images of centrioles in mitosis treated with DMSO, high-dose nocodazole (10µM) or taxol for 3 hours. Scale bar: 200 nm. **(i)** Quantification of the frequency of the centrosomes, in which the centrioles are connected at obtuse angle in (h). n = 40 centrioles pooled from 3 independent experiments. P values were calculated by Dunnett’s multiple comparisons test. **(j)** Representative immunofluorescence images of HeLa cells in mitosis transfected with siControl or siPCNT, in the presence or absence of high-dose nocodazole or taxol. **(k)** Quantification of the frequency of mitotic cells with disengaged centrioles in (j). n = 3 independent experiments, 30 cells each. P values were calculated by Dunnett’s multiple comparisons test. **(l)** STED microscopy images of centrioles in G2 phase. Scale bar: 200 nm. **(m)** Quantification of the locations of daughter centrioles and centrosomal proteins in regard to the center of the mother centriole. The end of the signal was defined as the location where the signal intensity dropped to 20% of its peak value. n = 30 centrioles pooled from 3 independent experiments. **(n)** Schematic model for the release of the torus engagement pathway by daughter centriole distancing. Data are represented as mean ± s.d.

A closer look at the centriole wall before and after centriole distancing revealed that the α -tubulin signal of the daughter’s proximal end is located farther from the mother centriole in mitosis, compared to G2 (Fig. 5e). In contrast, its acetylated tubulin signal, a marker of stabilized microtubules, appeared distant from the mother centriole already in G2 (Fig. 5e, f, g). Interestingly, the gap between the daughter’s proximal end signals of acetylated tubulin and α-tubulin observed in G2 phase was lost in mitosis (Fig. s5a, b, c). This is concomitant with the disappearing of the cartwheel structure during the transition from S/G2 phase to mitosis (Stage 4) ^38^. These observations suggest that the non-acetylated microtubules present at the base of the daughter centriole in G2 phase are lost as the cell progresses into mitosis. Given that the PLK1 inhibitor blocks both cartwheel removal and centriole distancing ^9,37^, we hypothesized that the process of centriole distancing is initiated by the loss of the daughter centriole’s proximal end.

To determine whether the loss of the proximal end is attributed to microtubule depolymerization from the minus-end, we treated cells with a high concentration (10 μM) of nocodazole. Although nocodazole is typically used as a microtubule-destabilizing agent, high concentrations of this drug can stabilize microtubule minus-ends ^39^. Cells treated with high-concentration nocodazole showed a higher fraction of centrioles associated under an orthogonal angle in mitosis compared to control cells (Fig. 5h, i, >100° centrioles, DMSO: 49.9±2.5%, Nocodazole: 23.5±4.2%). In contrast, treatment with taxol, which stabilizes the plus-but not the minus-end ^40^, did not suppress the obtuse angle formation of centrioles in mitosis (Fig. 5h, i, 55.5±4.5%). High-concentration nocodazole treatment also suppressed cartwheel detachment in mitosis (Fig. s5d, e, STLC: 18.9±6.9%, Nocodazole: 48.8±15.7%). Altogether, these data suggest that the loss of the daughter centriole’s proximal end is induced by microtubule depolymerization, which subsequently leads to centriole distancing.

We next investigated whether the mechanism of centriole engagement changes from Stage 3 to Stage 4 due to the centriole distancing by treating cells with high-concentration nocodazole. This treatment blocked the precocious centriole disengagement in mitosis caused by disturbing the PCM pathway with PCNT depletion (Fig. 5j, k, 10.0 ± 5.8%). In contrast treatment with taxol or low-concentration nocodazole had no such effect (Fig. 5j, k, s6a, b). Furthermore, CPAP depletion, which has been shown to inhibit centriole distancing ^37^, also suppressed precocious centriole disengagement in mitosis under depletion of Cep57 or PCNT (Fig. s6c, d). These results suggest that the centriole distancing mediated by the loss of the proximal end of the daughter centriole facilitates the transition of the centriole engagement mechanism from Stage 3 to Stage 4.

The alpha-tubulin signal at the proximal end overlapped with the ring of Cep63, whereas the acetylated tubulin signal, presumably corresponding to the mitotic daughter centriole, did not (Fig. 5l, m, n). In contrast, both signals overlapped with the ring of PCNT (Fig. 5f, l, m, n). These observations suggest that the daughter centriole in the G2 phase contacts both the torus and the PCM, while the torus is separated in mitosis due to centriole distancing. This change in the configuration corresponds to the change in the mechanism of centriole engagement, whereby the PCM pathway alone becomes responsible for the engagement in mitosis. We therefore propose that centriole distancing changes the engagement mechanism by releasing the daughter centriole from the torus engagement pathway, leading to the transition from Stage 3 to Stage 4.

### The stepwise changes in centriole engagement are necessary for proper centriole disengagement and centrosome number control

Our findings reveal that centriole engagement experiences unique alterations at different timings of the cell cycle. To determine whether these alterations occur in a stepwise manner, with each change depending on the completion of the previous one, we observed the consequences of halting each transition of centriole engagement. In control cells, a narrow daughter centriole is formed at the wall of its mother at an unstable angle, and then undergoes orthogonal adjustment (early S), blooming (late S), distancing with obtuse angle displacement (mitotic entry), and disengagement (after cell division) as the cell cycle progresses (Fig. 1a). When the stabilization of the orthogonal engagement angle in early S phase was inhibited by depleting cyclin E, daughter centriole blooming (late S) did not occur even as the cell cycle progressed leaving the centrioles in a thin unstable angle state (Fig. 6a). Similarly, when daughter centriole blooming (late S) was inhibited by depleting cyclin A2, the obtuse angle formation (mitotic entry) did not occur in mitosis (Fig. 6a). Furthermore, the failure in centriole blooming (late S) also resulted in a defect in centriole disengagement (after cell division) (Fig. 6a, b, c). Treatment with nocodazole or with the PLK1 inhibitor, which blocks centriole distancing (mitotic entry), also prevented centriole disengagement (after cell division) (Fig. 6a, b, c). Therefore, the configuration of the centriole engagement undergoes stepwise transitions throughout the cell cycle, and each of these is necessary for the eventual centriole disengagement.

**Figure 6.**
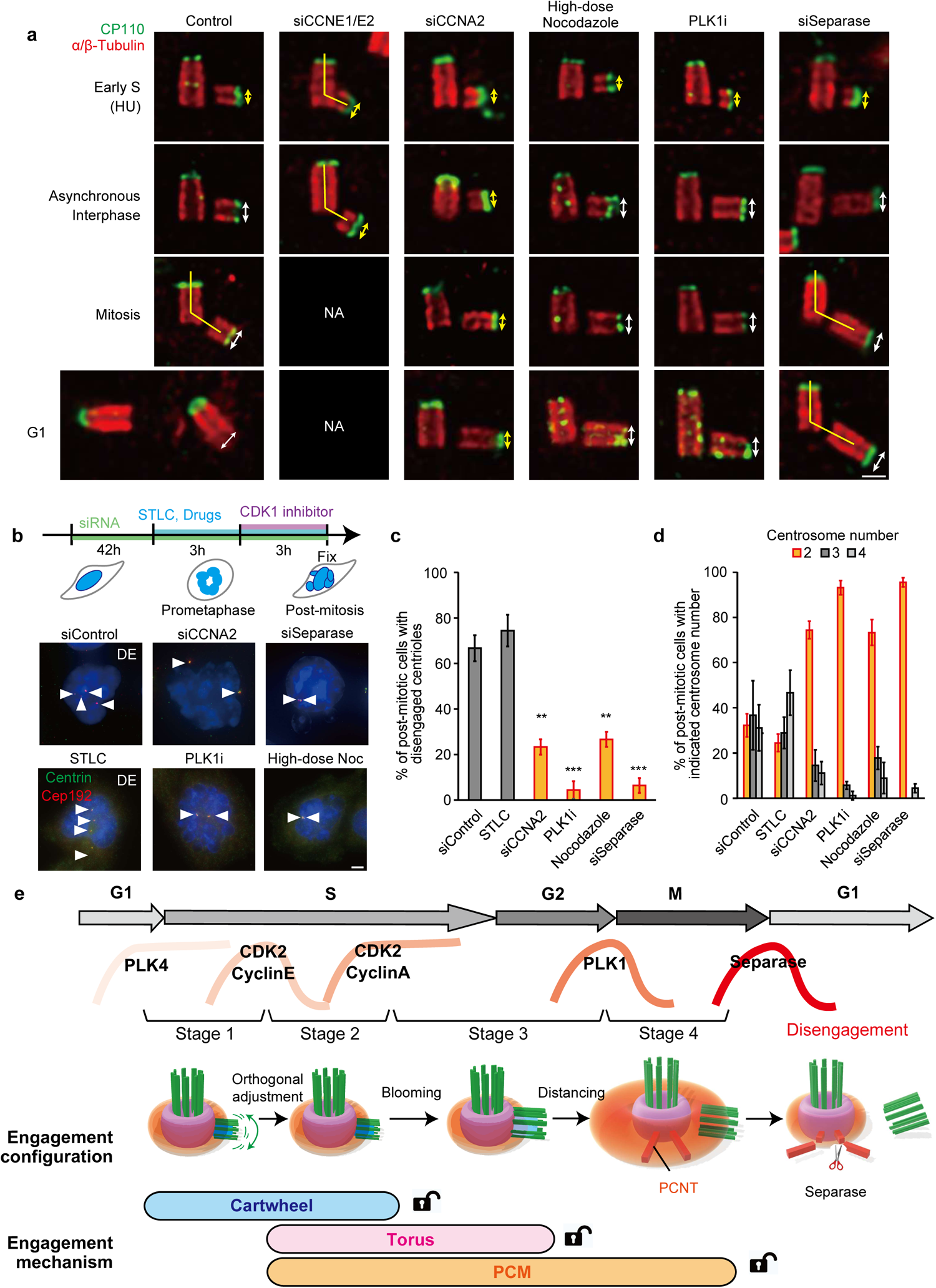
The stepwise changes of the centriole engagement are critical for daughter centriole release and its centriole-to-centrosome conversion. **(a)** U-ExM images of centrioles in HeLa cells treated with indicated siRNAs (48 h) or drugs (3 h). The G1 images were taken after the treatment shown in (b). Scale bars: 200 nm. Yellow arrows indicate the narrow centrioles. **(b)** Representative immunofluorescence images of HeLa cells post-mitosis upon the treatments with indicated siRNAs or drugs. **(c)** Quantification of the frequency of post-mitotic cells with disengaged centrioles in (b). n = 3 independent experiments, 30 cells each. P values were calculated by Dunnett’s multiple comparisons test. **(d)** Quantification of the frequency of the indicated centrosome numbers determined through the number of Cep192 foci in (b). n = 3 independent experiments, 30 cells each. **(e)** Schematic model of the four stages of centriole engagement throughout the cell cycle. Each stage is characterized by a distinct mechanism and a unique configuration of the centriole engagement. The changes in the engagement are caused by the structural maturation of the daughter centriole. Data are represented as mean ± s.d.

The disengaged daughter centriole becomes an independent, functional centrosome in the next G1 phase. We therefore investigated the effect of inhibiting the stepwise engagement changes on the conversion process of centrioles to centrosomes. Because some but not all treatments that block the changes in centriole engagement also caused mitotic arrest, we first decided to supplement STLC to all conditions in order to arrest all cells in mitosis, and then to force them into G1 phase by adding a Cdk1 inhibitor ^41^. Since cytokinesis fails under this condition, the post-mitotic cells inherit both centrosomes that formed the mitotic spindle pole, with each of them containing a mother and a daughter centriole. In the subsequent G1 phase, these centrioles are disengaged and the independent daughters are converted into centrosomes. As a result, the cell ends up with more than two centrosomes in the control condition (Fig. 6b, d). In contrast, inhibiting any of the changes in centriole engagement reduced the number of converted centrosomes (Fig. 6b, d). These findings indicate that the stepwise changes in the centriole engagement are essential for the ultimate disengagement of centrioles, enabling the daughter centriole to convert into a functional centrosome (Fig. 6e).

## Discussion

Centriole engagement is strictly maintained during interphase but swiftly disrupted following mitosis for centrosome number control. However, the mechanisms responsible for this accurate temporal control were unclear. Our study revealed that the centriole engagement is maintained strictly through three redundant pathways—the cartwheel, the torus, and the PCM pathways (Fig. 6e). The cartwheel and the torus are sequentially released before mitosis by two distinct steps of the progressive maturation of the daughter centriole: the centriole blooming and distancing, respectively. The release of these engagement pathways allows for the swift disruption of centriole engagement after mitosis, which is accomplished through the disassembly of the remaining PCM pathway. Thus, we showed that the daughter centriole gradually matures concomitant with the advance of the cell cycle to ultimately break the engagement with its mother and become an independent centrosome.

Prior to the stable tethering that occurs in Stage 2, the newly formed daughter centriole relies on the cartwheel for its initial engagement to the mother centriole (Stage 1). During Stage 1, the link between the daughter and the mother centrioles is flexible, as indicated by the variable degrees of engagement angles observed. The daughter centriole is rapidly broken by treatments inducing precocious disengagement, implying its immaturity. This property is thought to play a role for limiting the number of centrioles in duplication. In the case of multiple potential duplication sites arising simultaneously, the final duplication site is ultimately restricted to a single location ^42^. It is assumed that the immaturity of the daughter centrioles enables the removal of any extra procentrioles to establish only one. Later on, when the daughter centriole reaches Stage 2, it becomes connected to the mother centriole at a stable angle. Moreover, the Stage 2 daughter centriole is stable enough to retain its structure even after induction of precocious disengagement. Interestingly, we found that the transition from Stage 1 to Stage 2 is driven by the activation of the CDK2-cyclinE complex, which occurs at the restriction point ^43^. The restriction point is a critical checkpoint in the cell cycle, after which the cell is committed to dividing and cannot return to a quiescent state ^43^. Therefore, the restriction point works as a point-of-no-return not just for the cell cycle progression but also for daughter centriole formation.

After the restriction point, the centrioles are orthogonally engaged. The orthogonal positioning of the daughter centriole in respect to the mother has been observed in almost all animals ^44,45,46^. We have now discovered for the first time that this orthogonal configuration of centriole engagement is established by the torus protein Cep63 in human cells. Interestingly, the torus proteins, such as Cep63, Cep57 and Cep57L1 lack homologs in nematodes and insects. In addition, several structural differences between the centrioles of nematodes and insects, and those of humans have been reported ^45,46^. For human centrioles the blooming occurs as the diameter expands, accompanied by the acquisition of triplet microtubules ^7^. During this process, the cartwheel loses its contribution to the engagement as the distance between the cartwheel and the inner wall of the daughter centriole increases, before it ultimately dissociates during mitosis. In contrast, in nematode and insect centrioles, which only possess singlet and doublet microtubules, respectively, the cartwheel remains associated even beyond mitosis ^45,46^. Therefore, it is possible that the cartwheel maintains the engagement throughout interphase in these organisms. Intriguingly, orthogonal engagement is established in these species as well, suggesting that regardless of the underlying mechanism, the orthogonal connection represents a robust and fundamental aspect of centriole engagement. In the field of construction, orthogonal structures are well-suited for withstanding external pressures and forces ^47^. Thus, the orthogonal configuration appears to be a fundamental and conserved feature of centriole engagement across various species.

Numerical abnormalities in centrosomes are reported to lead to cancer progression and malignancy ^48^. In cancer cells, structural abnormalities are also observed ^49^. In this study, we demonstrated that the stepwise structural maturation of the daughter centriole leads to changes in centriole engagement. Disruptions in any transition between the stages of this maturation can contribute to precocious centriole disengagement and centrosome amplification. Among the important molecules related to these transitions, dysregulation of Cep63, cyclin E, cyclin A, and PLK1 are reported to lead to the accumulation of abnormal numbers of centrosomes and are thus included in the list of CA20, a set of genes used as an index for centrosome amplification in cancer cells^50^. Our findings suggest that the precise regulation of the stepwise structural maturation of the daughter centrioles may be critical to prevent centrosome amplification in cancer cells. Overall, we provide a comprehensive overview of the processes regulating mother-daughter centriole engagement throughout all phases of the cell cycle. We elucidate the three pathways underlying the strict temporary transitions between the maintenance and the stepwise disruption of centriole engagement, which likely play a key role in the control of centrosome numbers in cancer. These findings contribute novel insights for the development of therapeutic strategies.

## Figure legends

**Supplementary Figure 1 Cep57 and Cep57L1 maintain centriole engagement in interphase (a)** Representative immunofluorescence images of HeLa cells in early S phase, late S phase, G2 phase and mitosis. Scale bar: 200 nm. **(b)** Quantification of the frequency of cells with disengaged centrioles in G2 phase in (a). n = 3 independent experiments, 50 cells each. P value was calculated by Dunnett’s multiple comparisons test. **(c)** Quantification of the frequency of cells with disengaged centrioles in mitosis in (a). n = 3 independent experiments, 30 cells each. P value was calculated by Dunnett’s multiple comparisons test. Data are represented as mean ± s.d.

**Supplementary Figure 2 Confirmation of the centriole engagement by Cep57 and Cep57L1 in interphase using knockout cell lines (a)** Schematic illustration for generating knockout (KO) cells using CRISPR-Cas9. **(b)** immunofluorescence images of HeLa-Cas9, Cep57KO and Cep57L1KO cells. Scale bar: 200 nm. **(c)** DNA sequences surrounding the CRISPR-targeted regions in the exons of the Cep57 and Cep57L1 genes. **(d)** Representative immunofluorescence images of HeLa-Cep57KO cells transfected with siCep57L1, and Cep57L1KO cells transfected with siCep57 in G2 phase. Scale bar: 200 nm. **(e)** Quantification of the frequency of cells possessing four disengaged centrioles and more than four centrioles in G2 phase in (d). n = 3 independent experiments, 50 cells each. P value was calculated by two-tailed unpaired Welch’s t-test. **(f)** Representative immunofluorescence images of HeLa-Cep57L1KO cells transfected with siCep57L1 in early and late S phase. Scale bar: 200 nm. **(g)** Quantification of the frequency of the cells with disengaged centrioles in early S, late S and G2 phase in (f). n = 3 independent experiments, 30 cells each. P values were calculated by Dunnett’s multiple comparisons test. **(h)** Representative immunofluorescence images of HeLa cells subjected to a cartwheel removal assay. HeLa cells were treated with HU and Cdk2 inhibitor III (Cdk2i) for 24 hours, followed by treatment with a PLK4 inhibitor for another 24 hours. Scale bar: 5 µm. Data are represented as mean ± s.d.

**Supplementary Figure 3 Cep63 is involved in centriole engagement (a)** Immunofluorescence images of HeLa-Cas9 and Cep63KO cells in interphase. Scale bar: 200 nm **(b)** Immunofluorescence images of HeLa-Cas9 and Cep63KO cells in mitosis. Scale bar: 200 nm **(c)** DNA sequences surrounding the CRISPR-targeted regions in the exons of the *Cep63* gene. **(d)** Quantification of the frequency of mitotic cells with the indicated spindle shape. **(e)** Quantification of the frequency of mitotic cells with indicated number of centrioles in (b). **(f)** Representative immunofluorescence images of HeLa-Cas9 and Cep63KO cells transfected with siControl, siCep57 or siPCNT in interphase. Scale bar: 200 nm. **(g)** Quantification of the frequency of cells with disengaged centrioles in (f). P values were calculated by Dunnett’s multiple comparisons test. **(h)** Immunofluorescence images of HeLa cells overexpressing Cep57, Cep57L1, Cep63, or PCNT after plasmid transfection. P value was calculated by two-tailed unpaired Welch’s t-test. **(i)** STED microscopy image showing the top view of a mother centriole in G2 and mitosis **(j)** STED microscopy image showing the side view of a mother centriole in G2 and mitosis. Data are represented as mean ± s.d.

**Supplementary Figure 4 PLK1 overexpression changes the configuration and the mechanism of centriole engagement** (a) Representative STED microscopy images of HeLa-tet3G-PLK1T210D cells treated with DMSO or Doxycycline in HU-arrested S phase. Scale bar: 200 nm. (b) Quantification of the frequency of cells with indicated centriole angle in (a). (c) Representative immunofluorescence images of HeLa-tet3G-PLK1T210D cells treated with DMSO or Doxycycline in HU-arrested S phase. Scale bar: 5 μm. (d) Quantification of the frequency of cells with disengaged centrioles in (c). P values were calculated by Dunnett’s multiple comparisons test. Data are represented as mean ± s.d.

**Supplementary Figure 5 The proximal end of the daughter centriole is depolymerized in mitosis (a)** Representative STED microscopy images of centrioles in HeLa cells in G2 phase or mitosis. Scale bar: 200 nm. **(b)** Quantification of the signal intensity of α- and acetylated tubulin in (a). **(c)** Quantification of the daughter centrioles which had non-acetylated tubulin at their proximal ends in (a). The distance between mother centriole wall and daughter proximal end was determined by measuring the distance between the two points on either side of the signal valley where the intensity was 50% of the peak value. We defined a significant difference between the α-Tubulin and acetylated-Tubulin signals when the width of the valley differed by more than 20 nm. P value was calculated by two-tailed unpaired Welch’s t-test. **(d)** Representative immunofluorescence images of HeLa cells in mitosis treated with STLC, PLK1 inhibitor or high-dose nocodazole for 3 hours. Scale bar: 5 μm, 1 μm. **(e)** Quantification of the frequency of cells with cartwheel-retaining centrosomes in (d). P values were calculated by Dunnett’s multiple comparisons test. Data are represented as mean ± s.d.

**Supplementary figure 6 Centriole distancing is required for transition of engagement from Stage 3 to Stage 4 (a)** Representative immunofluorescence images of HeLa cells in mitosis transfected with siControl, siCep57 or siPCNT, under treatment with different doses of nocodazole. Scale bar: 5 μm, 1 μm. **(b)** Quantification of the frequency of cells with disengaged centrioles in mitosis in (a). P values were calculated by Dunnett’s multiple comparisons test. **(c)** Representative immunofluorescence images of HeLa cells transfected with siControl, siCep57 or siPCNT, in the presence or absence of siCPAP in mitosis. Scale bar: 5 μm. **(d)** Quantification of the frequency of cells with disengaged centrioles in (c). P value was calculated by two-tailed unpaired Welch’s t-test with Bonferroni correction. Data are represented as mean ± s.d.

## Methods

### Cell culture

HeLa cells were obtained from the European Collection of Authenticated Cell Cultures (ECACC). Cells were cultured in Dulbecco’s Modified Eagle’s Medium (DMEM) supplemented with 10% fetal bovine serum (FBS), 100 U/mL penicillin, and 100 μg/mL streptomycin at 37°C in a humidified 5% CO2 incubator.

### RNA interference

RNA interference experiments were performed by transfecting cells with Silencer Select siRNAs (Thermo Fisher) using Lipofectamine RNAiMAX (Life Technologies). The siRNAs used in this study were targeting the following transcripts: Cep57 (s18692) ^12^, Cep57L1 (s226224) ^14^, Cep63 (s37123) ^14^, PCNT (s10138) ^12^, CCNA2 (s2513), CCNE1 (s2524), CCNE2 (s17449), CEP295 (s229742) ^51^, CDK5RAP2 (s31430) ^12^, CPAP (s31623) ^51^, ESPL1 (s18686), and a scrambled siRNA was used as a negative control (4390843). The siRNAs for ESPL1, CCNA2, CCNE1, and CCNE2 were labeled as “Validated,” and their efficiency was confirmed by Thermo Fisher. The efficiency of the other siRNAs was confirmed in the references. All siRNAs were used at a final concentration of 20 nM and cells were treated for 48 hours unless otherwise specified.

### Antibodies

The following primary antibodies were used in this study: rabbit antibodies against Cep57 (GeneTex; GTX115931; IF, 1:1,000), Cep57L1 (Proteintech; 24957-1-AP; IF, 1:500), Cep63 (Proteintech; 16268-1-AP; IF, 1:1,000), PCNT (Abcam; ab4448; IF, 1:2,000), Cep192 (Bethyl Laboratories; A302-324A; IF, 1:1,000), CP110 (Proteintech; 12780-1-AP; IF, 1:500, U-ExM, 1:250), CENP-F (Abcam; ab108483; IF, 1:500), Ac-tubulin (Abcam; Ab179484; IF, 1:1,000); mouse antibodies against Cep57 (Abcam; ab169301; IF, 1:1,000), PCNT (Abcam; ab28144; IF, 1:1,000), centrin (Merck; 20H5; IF, 1:500), SASS6 (Santa Cruz; Sc-81431; IF, 1:300), α-tubulin (Merck; DM1A; IF, 1:1,000), Ac-tubulin (Sigma; T7451; IF, 1:1,000), CENPF (BD; 610768; IF, 1:500), CCNA2 (Cell Signaling; BF683; IF, 1:300), GTU88 (Merck; T5192; IF, 1:1,000); rat antibodies against PCNA (Abcam; ab252848; IF, 1:1,000), Centrin (Bio Legend; 698602; IF, 1:500); guinea pig antibodies against α-tubulin (ABCD antibodies; AA345; U-ExM, 1:250), β-tubulin (ABCD antibodies; AA344; U-ExM, 1:250), and goat antibody against PCNT (Santa Cruz; Sc-28145; IF, 1:200). Secondary antibodies used were donkey anti-mouse IgG (H+L) Alexa Fluor 488 (Invitrogen; A32766; IF, 1:500), donkey anti-rabbit IgG (H+L) Alexa Fluor 555 (Invitrogen; A31572; IF, 1:500), donkey anti-rat IgG (H+L) Alexa Fluor 647 (Invitrogen; A48272; IF, 1:500), camelid anti-rabbit IgG Abberior STAR 635P FluoTag-X2 (Abonva, RAB00968, IF, 1:500), llama/alpaca anti-mouse IgG-X2 Abberior Star 635P (Progen, 1A23, IF, 1:500), and goat anti-guinea pig IgG (H+L) Alexa Fluor 555 (Invitrogen, A21435; U-ExM, 1:250).

### Generation of knockout cell lines

To generate knockout cell lines, we utilized HeLa cells stably expressing Cas9 (HeLa-Cas9), which were previously established using a lentiviral system ^12^. Single guide RNAs (sgRNAs) were designed to target exons in the N-terminal region that are common to all isoforms of the target gene. The sgRNAs were synthesized by in vitro transcription from DNA oligonucleotide templates using the HiScribe T7 Transcription Kit (New England Biolabs). The transcribed products were subsequently purified using the RNA Clean & Concentrator (ZYMO RESEARCH). Purified sgRNAs were introduced into HeLa-Cas9 cells by transfection using Lipofectamine RNAiMAX (Life Technologies). Following transfection, single cell colonies were isolated by limiting dilution. The successful generation of the knockout cell lines was confirmed by the absence of immunofluorescence signal in microscopy and by sequencing of the targeted genomic region.

### Chemicals

The following chemicals were used in this study: RO3306 (SIGMA, SML0569, 10 µM), BI2536 (AdooQ, A10134, 200 nM), Hydroxyurea (Sigma, H8627, 2 mM), Centrinone (MedChem Express, HY-18682, 200 nM), CDK2 inhibitor III (Calbiochem, 238803, 200 μM), Nocodazole (Wako, 140-08531, 10 µM), Taxol (Tocris, 33069-62-4, 100 nM), and GSK461364 (MedChem Express, HY-50877, 200 nM). Unless otherwise specified, the PLK1 inhibitor referred to in the text is BI2536.

### Immunofluorescence staining (IF)

Cells cultured on 15 mm diameter, 0.12-0.17 mm thick round coverslips were fixed by immersion in methanol at −20°C for 7 minutes. The fixed cells were then washed three times with PBS and blocked for 30 minutes at room temperature in a PBS solution containing 1% BSA and 0.05% Triton X-100 (blocking buffer). Subsequently, the cells were incubated with primary antibodies diluted in blocking buffer for either 2 hours at room temperature or 12 hours at 4°C. After three washes with PBS, the cells were incubated with secondary antibodies in blocking buffer for 2 hours at room temperature. Finally, the cells were stained with Hoechst in PBS and mounted on glass slides using 90% glycerol.

For STED microscopy imaging of centrioles, cells were first placed at 4°C for 30 minutes to depolymerize cytoplasmic microtubules. The cells were then permeabilized by incubation with CSK buffer (25 mM HEPES pH 7.4, 50 mM NaCl, 1 mM EDTA, 3 mM MgCl2, 300 mM Sucrose, 0.5% Triton X-100) for 5 minutes prior to fixation.

### Microscopy

Fluorescence microscopy imaging and phenotype counting were performed using an Axioplan2 fluorescence microscope (Carl Zeiss) with 63x/1.4 NA plan-APOCHROMAT objectives. STED microscopy images were acquired using a Leica TCS SP8 STED 3X system with a Leica HC PLAPO 100x/1.40 oil STED WHITE objective, and a 660 nm gated STED laser.

### Ultrastructure expansion microscopy (U-ExM)

The U-ExM protocol was performed as described previously ^52^. Cells cultured on 15 mm diameter, 0.12-0.17 mm thick round coverslips were incubated in 2% acrylamide + 1.4% formaldehyde diluted in PBS for 3–12 h. The coverslips were then placed on 35 μL of monomer solution (19% sodium acrylate, 0.1% bis-acrylamide, and 10% acrylamide) with 0.5% TEMED and 0.5% APS for 5 min at 4°C and then for 1 h at 37°C. The gels were then incubated in the denaturation buffer (200mM SDS, 200mM NaCl, 50mM Tris pH 9) and boiled at 95°C for 90 min. Next, they were transferred into water at room temperature. After expansion, the gels were cut into quarters. The gels were then incubated with primary antibodies diluted in a blocking buffer for more than 3 hours at 37°C. After three washes with PBS, the cells were incubated with secondary antibodies in the blocking buffer for 2 hours at 37°C.

## Acknowledgements

22H02629, 22K20624, 23K14176, 23H02627, 24K02174) from the Ministry of Education, Pharmaceutical Sciences.

## Author contributions

K.K.I, K.M., S.H., and D.K. designed the study. K.K.I., K.T., and K.N. performed experiments. K.K.I., K.M., S.H., and D.K. analyzed data. M.F., S.Y., and T.C. provided suggestions. K.K.I., S.H., and D.K. wrote the manuscript. All authors contributed to discussions and manuscript preparation.

## Competing interests

The authors declare no competing interests.

